# RabbitCD-HIT: Distributed High-Sensitivity Greedy Incremental Clustering for Billion-Scale Protein Sequences in Hours

**DOI:** 10.64898/2026.11.05.764756

**Authors:** Yijie Gao, Guiliang Ma, Zekun Yin, Lifeng Yan, Yang Yang, Tong Zhang, Tiancheng Zhao, Zhuo Liu, Xiaohui Duan, Beifang Niu, Bertil Schmidt, Weiguo Liu

## Abstract

We present RabbitCD-HIT, a scalable distributed high-sensitivity greedy incremental clustering tool for ultra-large protein sequence datasets. It fully exploits cluster resources to enable efficient billion-scale clustering. RabbitCD-HIT clustered the March 10, 2026 NCBI nr snapshot (1.02 billion sequences; 487 GB FASTA) at 90% identity within 5 hours on 64 nodes; the resulting representatives and cluster assignments are released as a public database resource. On a 256-core server, RabbitCD-HIT achieved a 2.1× –6.6 ×speedup over CD-HIT with identical results under matched settings. Our approach addresses performance bottlenecks of traditional greedy incremental algorithms and provides an effective solution for large-scale protein sequence clustering.

## Background

Protein sequence clustering is widely used in many bioinformatics tasks, including redundancy removal, functional annotation, sequence searching, and the construction of nonredundant reference datasets [1]. By grouping similar sequences and selecting representative members for each cluster, protein sequence clustering reduces redundancy, streamlines downstream analyses, and improves computational efficiency [2, 3]. Clustering also helps reveal homologous relationships and organize protein families, thereby facilitating downstream functional inference and comparative analysis [4, 5, 6].

Increasing sequencing throughput, declining costs, and large-scale genome and metagenome projects have led to a surge in sequenced genomes and metagenomes [7, 8, 9, 10]. This rapid growth has driven the expansion of protein sequence databases and made large-scale clustering both increasingly important and increasingly challenging [11]. The NCBI non-redundant (nr) protein database has already exceeded 1 billion sequences as of March 10, 2026, and its growth continues to accelerate. Although existing clustering tools employ diverse algorithmic strategies, achieving both high efficiency and high accuracy at this scale remains challenging [12].

CD-HIT [13, 14] employs greedy incremental clustering with short-word (*k*-mer) filtering to reduce alignment costs. However, its in-memory data structures and single-node design cause memory consumption and runtime to increase rapidly as dataset size grows. UCLUST [15] follows a similar greedy paradigm, using heuristic anchoring and fast sequence search to accelerate clustering, but it also becomes memory-intensive and inefficient when applied to hundreds of millions of sequences. More recently, Linclust [12], implemented in the MMseqs2 suite [16], introduced an aggressive linear-time clustering strategy that enables extremely fast processing of massive protein datasets. Nevertheless, this high efficiency is achieved at the cost of reduced clustering sensitivity compared with greedy incremental pipelines. To improve sensitivity, Linclust/MMseqs2 [12] further provides cascaded clustering strategies, but these approaches are generally less efficient than Linclust alone.

Moreover, the above tools provide limited support for distributed computing platforms, which constrains their ability to scale beyond the resources of a single node. As datasets continue to grow, the lack of mature mechanisms for distributing data and workloads across multiple machines becomes a major bottleneck [17, 18]. Consequently, even algorithms with lower theoretical complexity may still struggle to simultaneously maintain efficiency, scalability, and clustering sensitivity on ultra-large datasets [19].

To address these challenges, we propose RabbitCD-HIT, a distributed extension of CD-HIT that reformulates greedy incremental clustering into a producer-consumer model for large-scale parallel execution. RabbitCD-HIT employs a hierarchical parallelization strategy that combines MPI-level process parallelism, OpenMP-level thread parallelism, and SIMD-level data parallelism. At the coarse-grained level, RabbitCD-HIT improves distributed and intra-node scalability through an MPI+OpenMP hybrid architecture, NUMA-aware execution, hybrid static and run-time load balancing, adaptive low-memory streaming I/O, and producer-consumer-aware process-thread allocation. At the fine-grained level, it accelerates compute-intensive kernels with SIMD vectorization, including a wavefront-based parallel alignment algorithm. These optimizations allow RabbitCD-HIT to efficiently exploit modern distributed clusters, substantially accelerate large-scale protein clustering, and preserve the clustering accuracy of the original CD-HIT.

On a server equipped with 256 CPU cores, RabbitCD-HIT achieves substantial speedup over the original CD-HIT, ranging from approximately 2.1× to 6.6× across the evaluated dataset sizes. On distributed clusters, RabbitCD-HIT further makes it practical to perform CD-HIT-compatible clustering on the full NCBI non-redundant protein database, completing the 90% sequence-identity clustering of approximately 1.02 billion protein sequences within only a few hours. Importantly, this computation also enables the construction of a publicly available 90% non-redundant nr resource, including the complete representative sequence set and full cluster assignment files, which we plan to update periodically as nr continues to expand. Together, these results demonstrate that RabbitCD-HIT not only improves the scalability of greedy incremental clustering, but also provides a reusable community resource for large-scale protein sequence analysis.

## Results

### Overview

RabbitCD-HIT is a distributed redesign of CD-HIT for ultra-large protein sequence clustering. CD-HIT follows a greedy incremental clustering strategy, in which input sequences are sorted by decreasing length and each sequence is assigned to an existing representative if it satisfies the short-word filtering and sequence identity criteria; otherwise, it is retained as a new representative. Owing to its sub-quadratic practical complexity and high clustering sensitivity, CD-HIT has been widely adopted by the bioinformatics community and has served as a de facto standard tool for protein sequence clustering over the past two decades. However, the original CD-HIT implementation lacks native support for distributed computing, which limits its ability to exploit large-scale clusters and makes it difficult to process ultra-large protein databases containing billions or even larger amount of sequences.

RabbitCD-HIT produces identical clustering results to CD-HIT while parallelizing the expensive sequence-checking procedure across distributed computing nodes. As illustrated in Figure 1, RabbitCD-HIT partitions the sorted input sequences into chunks and adopts a producer–consumer execution model. The master (producer) process selects representative sequences according to the global greedy order and broadcasts compact representative metadata to worker (consumer) processes. Each worker then constructs local short-word inverted indices and checks its assigned sequences against the received representatives using the same filtering and alignment rules as CD-HIT. The local clustering results are finally returned to the master and merged into the final output. Under matched clustering settings, RabbitCD-HIT produced clustering assignments fully consistent with those of the original CD-HIT, confirming that the distributed execution does not alter CD-HIT-compatible clustering semantics.

**Figure 1.**
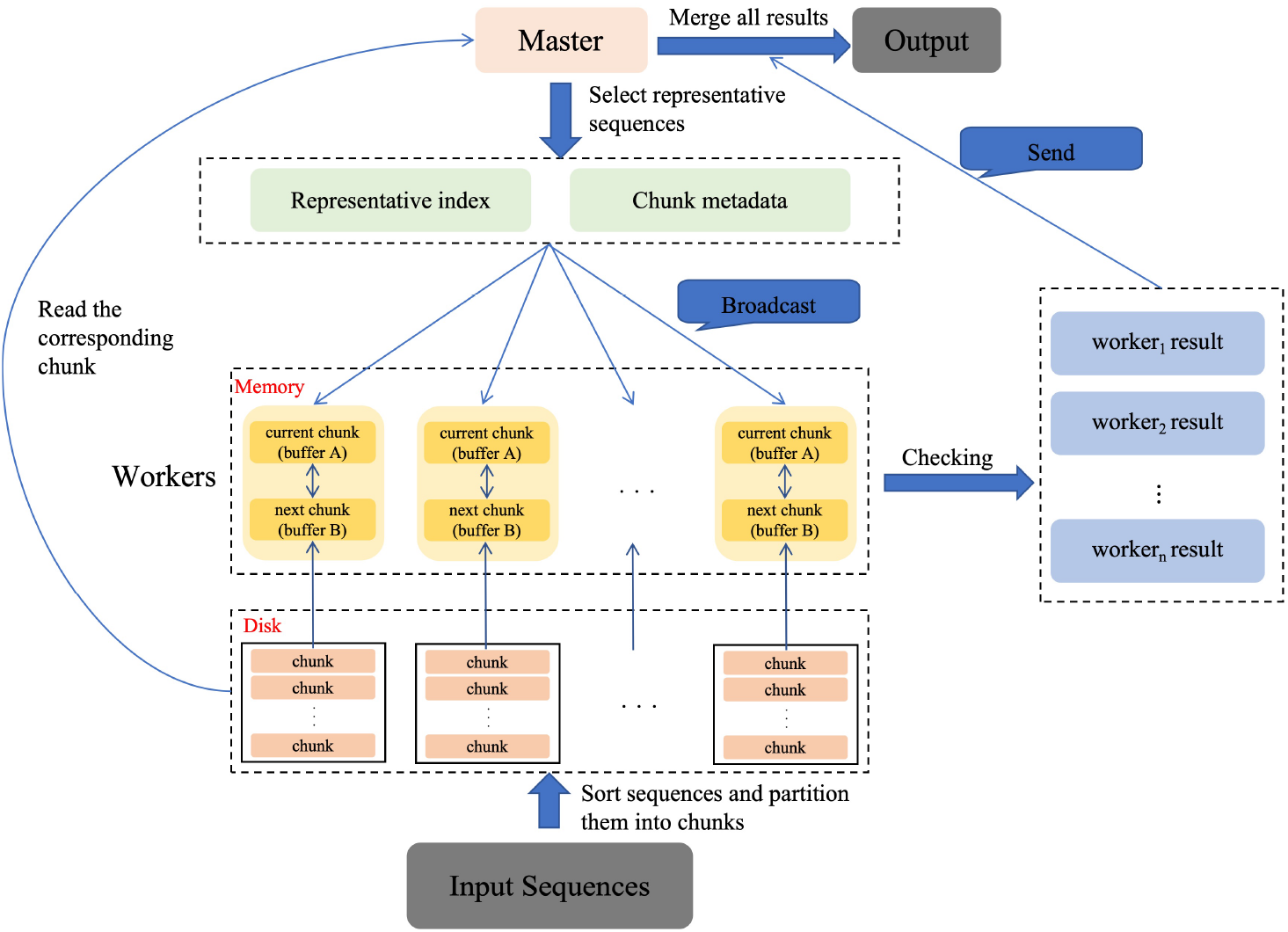
Overview of RabbitCD-HIT.

To support memory-efficient execution, RabbitCD-HIT adopts an on-demand I/O strategy with double-buffered prefetching, where sequence chunks are loaded only when needed and the next required chunk is prefetched into a second worker-local buffer while the current chunk is being checked, thereby avoiding full database loading and overlapping disk I/O with computation.

### Clustering 1.02 Billion Protein Sequences within Hours

We applied RabbitCD-HIT to the NCBI non-redundant protein sequence database snapshot released on March 10, 2026. This dataset contains approximately 1.02 billion protein sequences and occupies 487 GB in FASTA format. RabbitCD-HIT clustered the full dataset at a 90% sequence identity threshold on 64 compute nodes. Each node was equipped with two AMD EPYC 9755 processors, providing 256 physical CPU cores per node.

RabbitCD-HIT completed the full clustering task in 257 minutes, whereas CD-HIT did not finish the same task within one week. The clustering results contained 415,493,515 clusters and the largest cluster comprised 1,271,468 sequences. This corresponds to an average processing throughput of approximately 4.0 million sequences per minute, or about 66,000 sequences per second. The resulting cluster count indicates that approximately 40.7% of the original sequences were retained as representatives at the 90% identity threshold, corresponding to an average cluster size of approximately 2.46 sequences per representative. This experiment demonstrates that RabbitCD-HIT makes CD-HIT-compatible greedy clustering practical for billion-scale protein sequence databases, a scale that is beyond the practical capability of the original single-node CD-HIT.

The resulting representative sequence set provides a ready-to-use 90% non-redundant protein sequence resource derived from the full NCBI nr database. Such a resource can be directly used for downstream protein sequence search, functional annotation, redundancy reduction, and large-scale comparative analyses. The representative sequence set and full cluster assignment files have been made publicly available, as described in the Availability of data and materials section, and will be updated periodically as nr continues to grow.

### Performance Comparison with Widely Used Protein Clustering Tools

Because the existing tools considered in this comparison are limited to single-node execution, we evaluated RabbitCD-HIT under the same single-node computational setting for fairness. We compared its runtime with CD-HIT (version 4.8.1), Linclust/MMseqs2 (Release 18), and Linclust (Release 18). The benchmark datasets were constructed from the same NCBI nr snapshot dated March 10, 2026, and contained 2 million, 10 million, 50 million, and 100 million protein sequences, corresponding to approximately 1 GB, 4.8 GB, 24 GB, and 48 GB of FASTA data, respectively. All experiments were performed on a dual-socket AMD EPYC 9755 server equipped with 256 physical CPU cores, AVX-512 support, and 768 GB of memory. Each tool was evaluated at sequence identity thresholds of 0.9 and 0.7 using 256 cores, with other parameters kept at their default settings.

As shown in Figure 2, RabbitCD-HIT consistently accelerated CD-HIT across all tested datasets and identity thresholds. Across the evaluated conditions, RabbitCD-HIT achieved approximately 2.1× to 6.6× speedup over the original CD-HIT. The speedup is mainly driven by RabbitCD-HIT’s NUMA-aware hybrid MPI+OpenMP design, which replaces heavily shared cross-NUMA data access with process-local inverted indices. This design improves memory locality and reduces memory-bandwidth contention, while optimized intra-chunk clustering and alignment kernels further contribute to the performance gain.

**Figure 2.**
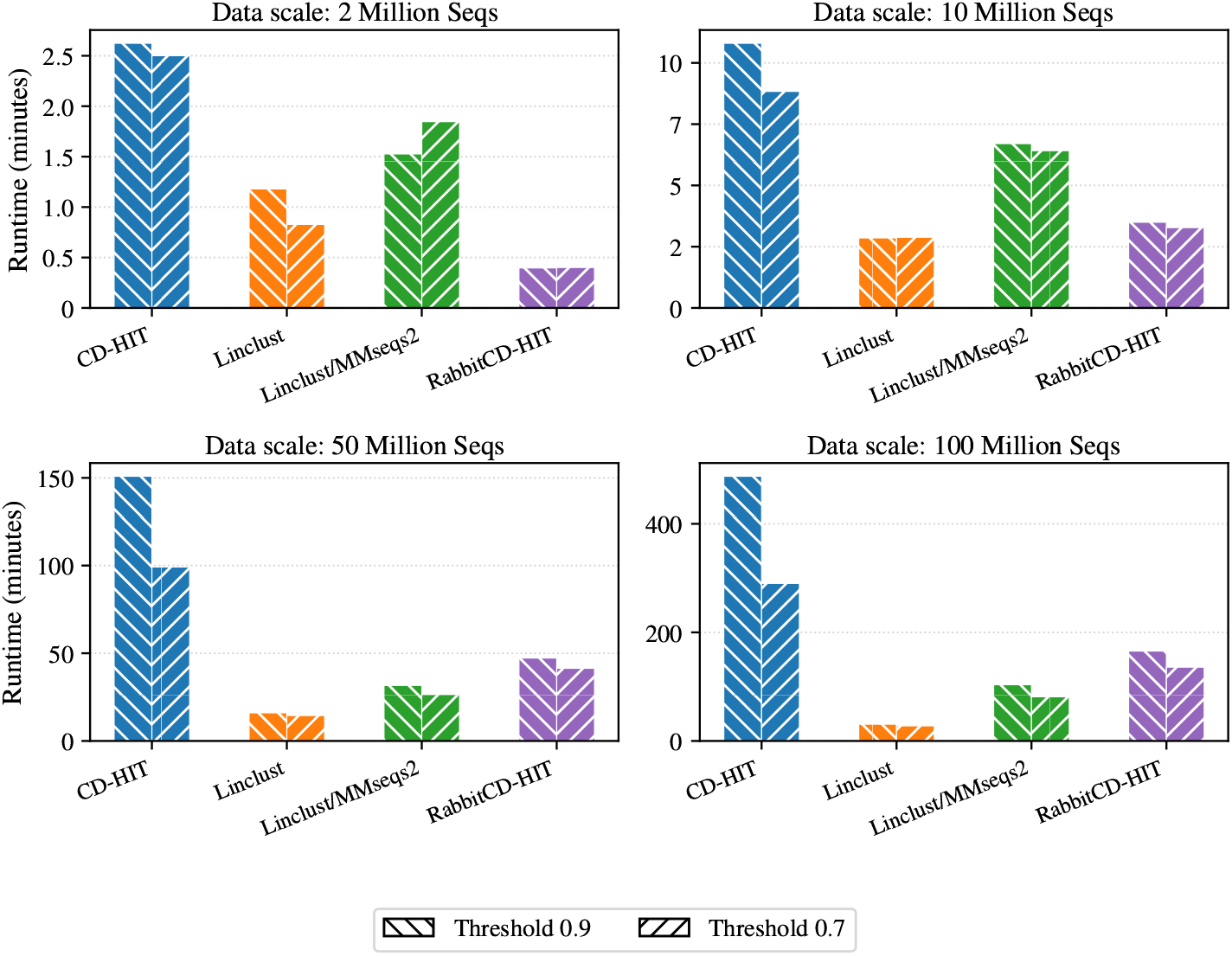
Runtime comparison of different clustering tools and data scales.

Compared with Linclust/MMseqs2, RabbitCD-HIT achieved lower runtime on the 2-million- and 10-million-sequence datasets under both sequence identity thresholds. On the 50-million- and 100-million-sequence datasets, Linclust/MMseqs2 achieved lower runtime, consistent with its highly optimized cascaded workflow that combines Linclust-based rapid initial clustering with multiple rounds of efficient cluster merging. Compared with Linclust, RabbitCD-HIT performed better on the smallest dataset and remained close on the 10-million-sequence dataset, whereas Linclust was substantially faster at larger scales because of its linear clustering strategy.

Although some existing tools can achieve lower runtime on a single node, they are fundamentally limited by single-node execution and lack mature support for exploiting distributed computing platforms. Consequently, their performance is bounded by the computational resources of a single server. RabbitCD-HIT overcomes this limitation by distributing sequence checking across multiple nodes, thereby leveraging cluster-scale computational resources to improve clustering throughput. This makes RabbitCD-HIT particularly advantageous for ultra-large protein databases, where the ability to scale across distributed platforms is more important than single-node speed alone.

We further compared the peak memory usage of different tools at a sequence identity threshold of 0.9 on the same single-node platform, using 256 CPU cores and otherwise default configurations. As shown in Table 1, RabbitCD-HIT used more memory than CD-HIT on smaller datasets, requiring 13.24 GB and 30.71 GB for the 2-million- and 10-million-sequence datasets, respectively. This additional memory usage is mainly introduced by the hybrid MPI+OpenMP execution model, where each MPI process maintains process-local inverted indices and auxiliary data structures for sequence checking. Such replicated structures introduce a near fixed per-process memory overhead, which is acceptable on modern multi-core servers.

**Table 1.** Single-node peak memory comparison of different clustering tools across data scales.

| Data scale | FASTA size (GB) | Peak memory usage (GB) |  |  |  |
|---|---|---|---|---|---|
|  |  | CD-HIT | RabbitCD-HIT | Linclust/MMseqs2 | Linclust |
| 2 million seqs. | 1.0 | 10.09 | 13.24 | 8.11 | **3.82** |
| 10 million seqs. | 4.8 | 21.03 | 30.71 | 24.42 | **15.95** |
| 50 million seqs. | 24.0 | **43.57** | 53.75 | 137.58 | 67.97 |
| 100 million seqs. | 48.0 | 71.04 | **69.77** | 145.28 | 135.75 |

However, as the dataset size increased, the memory growth of RabbitCD-HIT became much slower than that of the other tools. On the 50-million-sequence dataset, RabbitCD-HIT required 53.75 GB of memory, which was lower than Linclust/MMseqs2 and Linclust. On the 100-million-sequence dataset, RabbitCD-HIT used 69.77 GB, slightly lower than CD-HIT and substantially lower than Linclust/MMseqs2 and Linclust. This behavior is mainly due to the on-demand data loading strategy in RabbitCD-HIT, which avoids keeping the entire sequence database in memory. Instead, sequence chunks are loaded as needed during iterative checking, while the growth of auxiliary data structures remains relatively limited.

These results show that although RabbitCD-HIT introduces additional memory overhead on small datasets, its memory usage scales more robustly as the dataset grows. This design is particularly suitable for billion-scale and larger protein sequence databases. Moreover, RabbitCD-HIT is designed for distributed computing platforms, where the aggregate memory capacity increases with the number of nodes, making the per-process auxiliary memory overhead less significant at large scale.

### Strong Scalability on Distributed Computing Platforms

We further evaluated the distributed scalability of RabbitCD-HIT on the full nr workload. As shown in Figure 3, RabbitCD-HIT exhibited strong scaling from 1 node (256 cores) to 64 nodes (16,384 cores). Relative to the 1-node baseline, the measured speedups were 2.07×, 3.65×, 7.37×, 12.63×, 25.73×, and 35.43× on 2, 4, 8, 16, 32, and 64 nodes, respectively. The parallel efficiency remained approximately 80% at 32 nodes and approximately 55% at 64 nodes, indicating that RabbitCD-HIT can effectively exploit distributed computing resources for billion-scale clustering workloads.

**Figure 3.**
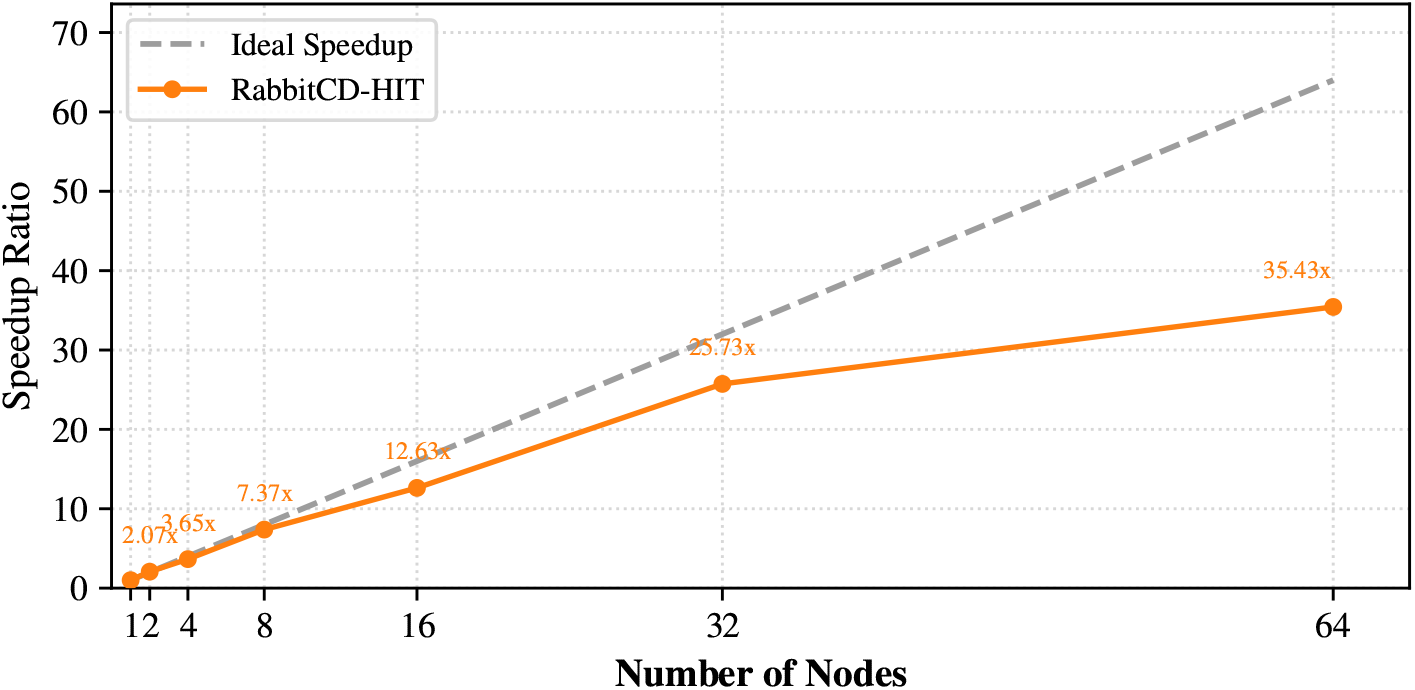
Distributed scaling of RabbitCD-HIT.

The speedup from 1 to 2 nodes was slightly superlinear. This behavior can be explained by the producer–consumer structure of RabbitCD-HIT. With 8 MPI processes per node, a 1-node configuration contains one producer process and seven worker processes, whereas a 2-node configuration contains one producer process and fifteen worker processes. Therefore, increasing from 1 to 2 nodes more than doubles the number of worker processes, improving the consumer-side processing rate.

From 2 to 32 nodes, the speedup increased nearly linearly, suggesting that chunk-level partitioning, distributed checking, and hybrid MPI+OpenMP parallelism effectively reduce per-node workload while keeping communication overhead manageable. At 64 nodes, the speedup continued to improve but deviated more clearly from the ideal linear trend. This is expected because the number of worker processes continues to increase, whereas the producer remains limited to a single process. As a result, representative selection, metadata preparation, and global synchronization on the producer side become increasingly important bottlenecks at larger scale.

Overall, the distributed scaling results show that RabbitCD-HIT can efficiently use multi-node computing resources and that its producer–consumer design is effective for scaling CD-HIT-compatible greedy clustering to billion-scale datasets.

### Performance Evaluation of System-level and Kernel-level Optimizations

RabbitCD-HIT combines several system-level and kernel-level optimizations to improve scalability and runtime efficiency. We evaluated two representative aspects of these optimizations: NUMA-aware execution on multi-socket servers and SIMD-accelerated banded alignment.

First, we examined the NUMA scalability of RabbitCD-HIT using the 10-million-sequence dataset. On the dual-socket AMD EPYC 9755 server with two NUMA domains, RabbitCD-HIT achieved a 1.88× speedup when scaling from one to two NUMA domains, whereas CD-HIT achieved only 1.20×, see Figure 4(a). This indicates that RabbitCD-HIT can more effectively exploit memory locality and parallel resources within a multi-socket server. We further evaluated NUMA behavior on a dual-socket AMD EPYC 7T83 server with eight NUMA domains. As shown in Figure 4(b), RabbitCD-HIT achieved approximately 2.09×, 4.4×, and 8.09× speedup when scaling to 2, 4, and 8 NUMA domains, respectively. In contrast, CD-HIT achieved only 1.37×, 1.89×, and 1.32× under the same configurations. These results show that RabbitCD-HIT has substantially better NUMA scalability than the original CD-HIT.

**Figure 4.**
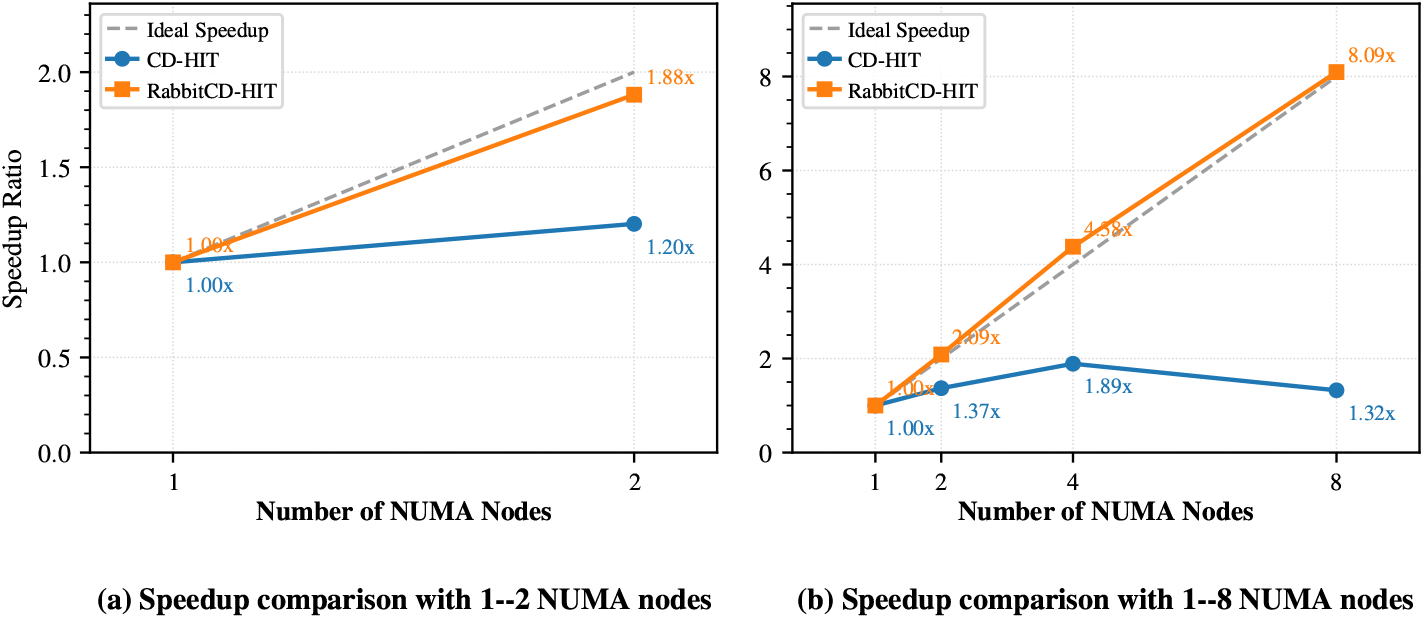
Speedup comparison with different NUMA node counts.

The improved NUMA scalability mainly comes from RabbitCD-HIT’s NUMA-aware MPI+OpenMP execution strategy. By binding MPI processes and OpenMP threads according to the NUMA topology, RabbitCD-HIT enables each process to maintain local inverted indices and process local data chunks whenever possible. Its process-thread allocation strategy further balances producer and consumer work-loads while preserving memory locality, thereby reducing remote memory traffic and memory-bandwidth contention compared with the original CD-HIT.

Second, we evaluated the optimized banded alignment kernel used after short-word filtering. We compared three implementations: the original row-major dynamic programming formulation, a compact wavefront formulation in rotated coordinates with no vectorization, and an AVX-512 vectorized implementation based on the compact wavefront layout. The results are shown in Figure 5. The compact wave-front formulation alone was slower than the original row-major implementation, with runtimes increasing by 1.31–3.27× across the tested sequence lengths, because coordinate rotation introduces additional state management and arithmetic over-head. However, this transformation removes within-row loop-carried dependencies and stores active cells in a more regular layout, making the computation suitable for SIMD vectorization. With AVX-512 vectorization, the optimized alignment kernel achieved clear performance improvements, reducing runtime by 1.46–2.17× compared with the original implementation and by 2.61–4.78× compared with the compact scalar formulation. The improvement was especially evident for longer protein sequences. As sequence length increases, the alignment workload becomes more compute-intensive, allowing the vectorized implementation to amortize the over-head of the rotated coordinate layout and evaluate multiple dynamic programming states in parallel. Therefore, the wavefront formulation should be viewed as an enabling transformation: it may not reduce scalar runtime by itself, but it creates the regular dependency structure needed for efficient SIMD execution.

**Figure 5.**
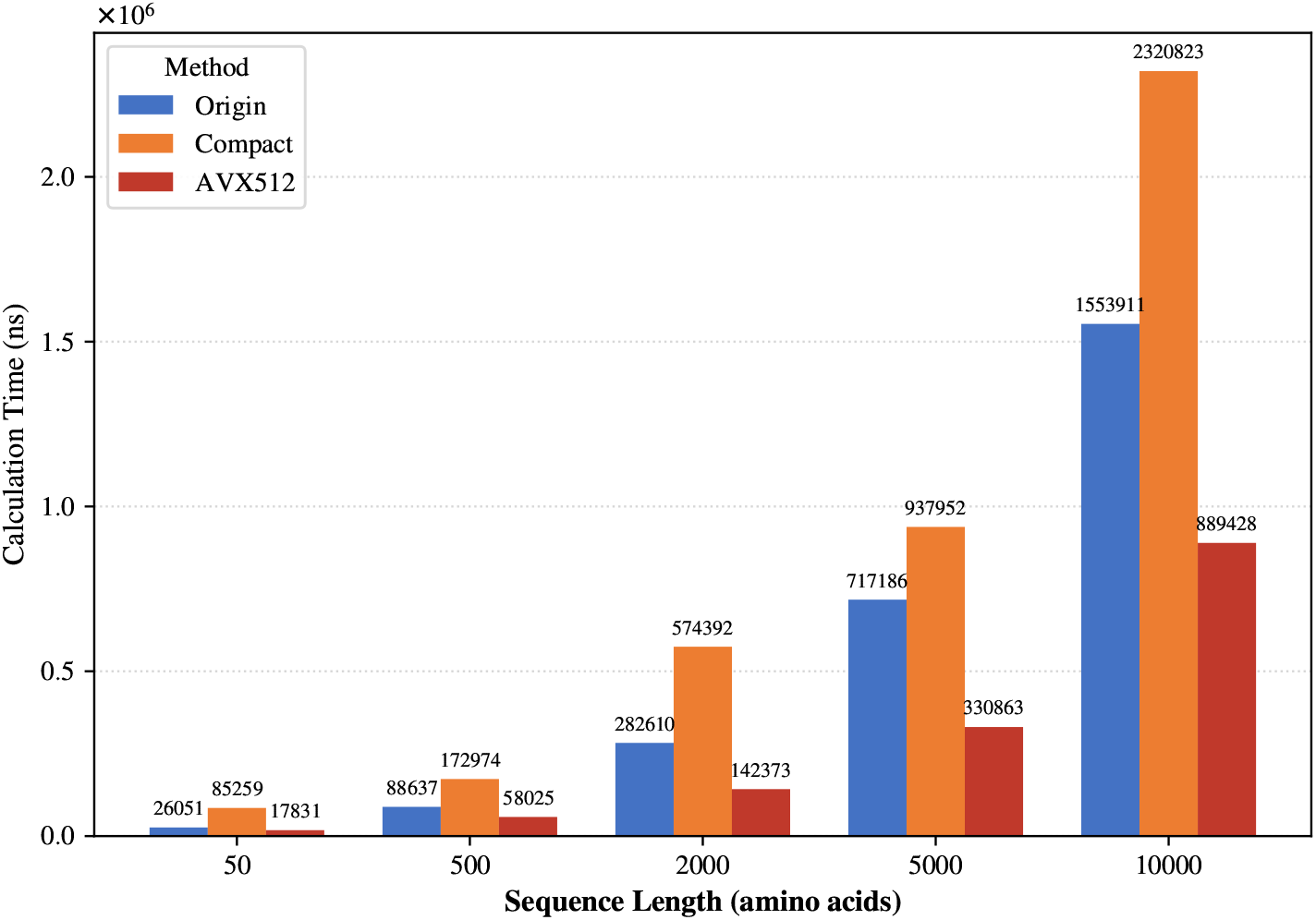
The local alignment performance comparison with different length.

Together, these results show that RabbitCD-HIT’s performance improvement comes from both system-level scalability and kernel-level acceleration. NUMA-aware process-level decomposition improves memory access locality and within-node scaling, while the vectorized banded alignment kernel reduces the cost of expensive sequence alignment. Combined with other optimizations detailed in Methods section, these optimizations enable RabbitCD-HIT to process billion-scale protein sequence datasets efficiently.

## Discussion and Conclusions

Large-scale protein sequence clustering requires a careful balance between computational efficiency, clustering sensitivity, and practical scalability [20]. Existing tools address this problem from different perspectives. Methods such as Linclust achieve very high throughput by adopting linear clustering strategies, making them particularly effective for rapid large-scale redundancy reduction. Greedy incremental approaches such as CD-HIT, on the other hand, provide well-defined clustering decisions based on short-word filtering followed by sequence alignment, but their practical scalability is constrained by single-node memory capacity and computational throughput.

RabbitCD-HIT is designed to extend this CD-HIT-style greedy clustering strategy to distributed computing platforms. By preserving the clustering semantics of CD-HIT while distributing data and workloads across multiple nodes, RabbitCD-HIT improves scalability without introducing additional approximation into the clustering procedure. As a result, RabbitCD-HIT offers a practical solution for high-sensitivity clustering of ultra-large protein sequence databases.

The main practical requirement of RabbitCD-HIT is access to distributed computing resources. Compared with single-node tools, this may introduce additional deployment and resource costs. However, such costs are often acceptable for large reference databases, which are typically clustered offline and updated periodically rather than continuously. Moreover, the resulting pre-clustered reference datasets can be shared and reused by the community, amortizing the computational cost across a broad range of downstream applications. RabbitCD-HIT therefore provides a scalable and reliable framework for constructing high-quality non-redundant protein sequence resources at billion-sequence scale.

## Methods

This section presents the design and implementation of RabbitCD-HIT. We first review the CD-HIT workflow and analyze its scalability limitations on modern multicore and NUMA-based systems. We then describe the overall RabbitCD-HIT architecture, including data preprocessing, chunk partitioning, and distributed producer-consumer execution. Finally, we introduce the key optimization techniques, including hierarchical MPI-OpenMP-SIMD parallelism, memory-efficient on-demand I/O, NUMA-aware process-thread allocation, and clustering consistency validation.

### Overview of CD-HIT

CD-HIT (Cluster Database at High Identity with Tolerance) is a widely used sequence clustering tool. It is based on the classical greedy incremental clustering algorithm that performs fast clustering by combining *k*-mer filtering with local sequence alignment.

The overall workflow of CD-HIT is illustrated in Figure 6. CD-HIT first sorts all input sequences from longest to shortest so that longer and more informative sequences are preferentially selected as cluster representatives. The clustering process follows a simple greedy strategy: each new sequence is compared only against the previously accepted representatives. CD-HIT processes the sequences chunk by chunk. CD-HIT first performs clustering within the current chunk (inner-clustering phase) to identify representative sequences. During the inner clustering stage, when a new sequence shows sufficient identity with an existing representative, it is assigned to that cluster; otherwise, it becomes a new representative. Then, the resulting representatives from the inner clustering are compared with the sequences in the remaining chunks to detect redundant sequences (checking phase).

**Figure 6.**
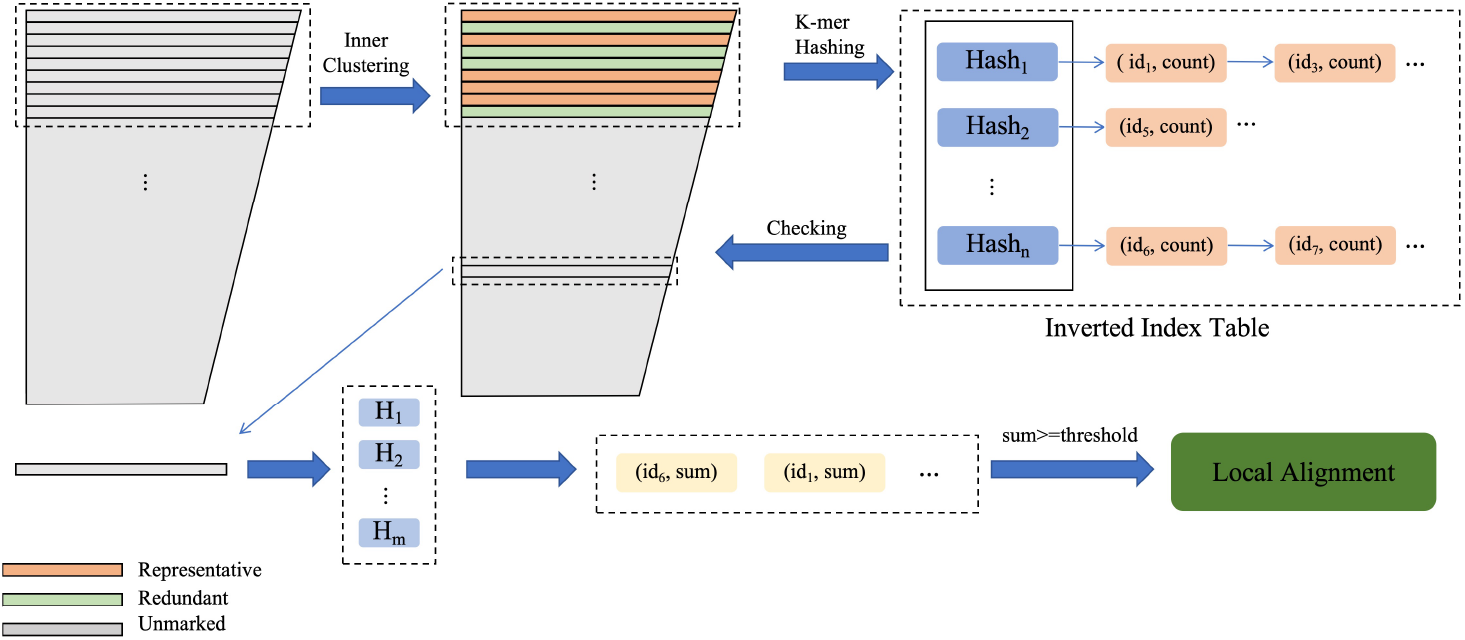
Overview of CD-HIT: (i) inner clustering: selects representative sequences within the current chunk and constructs a k-mer-based inverted index. (ii) checking: compares later sequences against these representatives through k-mer filtering and local alignment to identify redundant sequences.

To accelerate the clustering process, CD-HIT adopts a *k*-mer-based prefiltering strategy that substantially reduces the computational cost by eliminating unnecessary pairwise alignments. In this design, *k*-mer hits between sequences serve as a coarse similarity estimate. To efficiently detect such hits, CD-HIT constructs an inverted index of the *k*-mers belonging to the representative sequences identified during the inner-clustering stage. As illustrated in Figure 6, this index maps hashed *k*-mers (Hash_1_–Hash_*n*_) to lists of representative sequence IDs. When processing the remaining sequences, this structure enables rapid retrieval of candidates and efficient accumulation of their shared *k*-mer counts. Sequence pairs whose *k*-mer hit counts exceed a predefined threshold are treated as potential matches and passed to the more expensive alignment stage, whereas those falling below the threshold are immediately discarded. This inverted-index-based filtering mechanism substantially reduces the number of full alignments required and greatly improves overall performance.

CD-HIT supports OpenMP-based multithreading, which accelerates the checking phase on multi-core systems. However, this parallelization mainly benefits filtering and alignment operations, whereas the inner-clustering phase remains serial due to inherent data dependencies. While such a strategy is effective on conventional multi-core architectures, it may face scalability challenges on modern architectures with non-uniform memory access (NUMA) characteristics [21]. In modern multi-socket NUMA systems, memory is distributed across NUMA nodes, and remote memory accesses are typically more expensive than local accesses [22]. Since the inverted index is shared by all threads, it may lead to frequent remote memory accesses, thereby increasing memory latency, exacerbating bandwidth contention, and ultimately limiting multithreaded scalability.

Furthermore, efficient execution relies on keeping all sequence data and index structures entirely in main memory. As modern databases continue to expand, the NCBI non-redundant (nr) protein database already exceeds 450 GB as of March 10, 2026. Such memory demand can readily exceed the RAM capacity of typical commercial servers, leading either to out-of-memory failures or to severe disk swapping with prohibitive I/O overhead.

### Architecture of RabbitCD-HIT

RabbitCD-HIT is a distributed redesign of CD-HIT that combines MPI and OpenMP to enable large-scale sequence clustering on modern high-performance computing systems. We adopt this hybrid architecture to balance scalability, memory usage, and memory locality. A pure-MPI implementation would require each process to maintain its own copy of the inverted index, resulting in substantial memory duplication and limiting the practicality of clustering very large datasets. By contrast, a pure multithreaded implementation would avoid index replication, but on NUMA systems it could incur frequent remote memory accesses when many threads share the same data structure across memory domains. RabbitCD-HIT therefore uses MPI across processes and OpenMP within each process. This design is particularly important on multi-socket systems, where a socket may contain one or more NUMA nodes depending on the hardware topology; in our implementation, each process is restricted to at most one socket so that any unavoidable cross-NUMA accesses remain within the same socket, where the cost is typically lower than across sockets. On top of this hybrid parallel design, RabbitCD-HIT adopts a producer-consumer execution model, in which the producer (master) process performs inner clustering using a newly designed multithreaded method, while multiple consumer (worker) processes perform checking tasks in parallel, enabling efficient pipelined execution across the workflow. The workflow of RabbitCD-HIT is illustrated in Figure 7.

**Figure 7.**
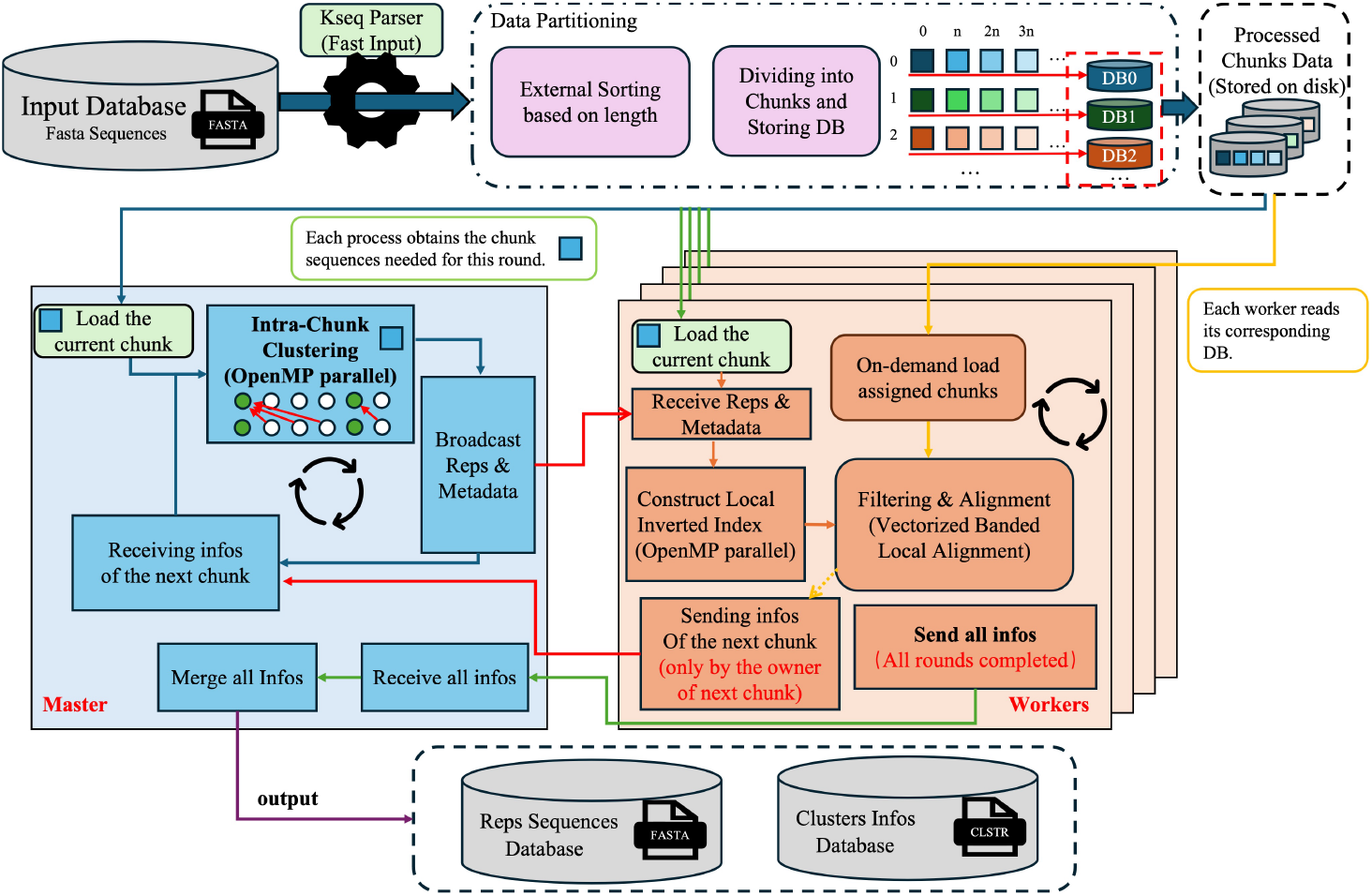
Workflow of RabbitCD-HIT. Master: performs inner clustering, broadcasts representative sequences and metadata to all worker processes for parallel checking. Workers: compare their assigned sequences against the received representative data and generate local clustering results. After all clustering rounds are completed, these results are sent back to the master process for merging and writing to disk.

RabbitCD-HIT consists of four main steps.

**Step 1**. The input database is first sorted, partitioned, and assigned to workers in a cyclic manner, providing a simple and deterministic static distribution that helps balance the workload. However, because the actual clustering cost depends on sequence similarity and runtime filtering behavior, static cyclic partitioning alone cannot guarantee perfect load balance. To address this issue, our framework adopts a double-buffered asynchronous pipeline, which enables neighboring chunks assigned to different workers to be processed asynchronously and overlaps communication operations, such as broadcasting, with checking computation. This design reduces worker idle time and improves overall parallel efficiency.

**Step 2**. The master process loads a chunk of sequences and performs efficient multithreaded inner-clustering to identify representative sequences. It then broad-casts the representative sequences in the current chunk, along with the necessary metadata, to all worker processes.

**Step 3**. Upon receiving the representative sequences and metadata, each worker process constructs the corresponding inverted index and performs the optimized checking operations. Meanwhile, the master process proceeds to load the next chunk and repeats **Step 2**, while the worker processes repeat **Step 3**. This workflow continues until all data chunks have been processed.

**Step 4**. After all clustering rounds have been completed, each worker process sends its final clustering results back to the master process. The master process then merges the received results and writes the consolidated output to disk.

### Data Preprocessing and Partitioning

To achieve balanced workload distribution, RabbitCD-HIT sorts and partitions the input protein sequences so that each consumer process receives a set of data chunks with a comparable total size.

Because the input dataset is typically very large and cannot be entirely loaded into memory for in-memory sorting, we design an efficient external sorting pipeline. The original FASTA file is first divided into multiple segments, which are sorted in parallel using multithreading to produce several ordered subfiles. These subfiles are then progressively merged using a merge-sorting strategy to generate a set of chunks with approximately equal sizes (the chunk size is estimated based on the total sequence length so that each chunk has an approximately equal total byte size), as illustrated in Figure 8(a). Each chunk is written to a separate file, allowing consumer processes to directly load and handle their assigned chunks without additional communication with the master process.

**Figure 8.**
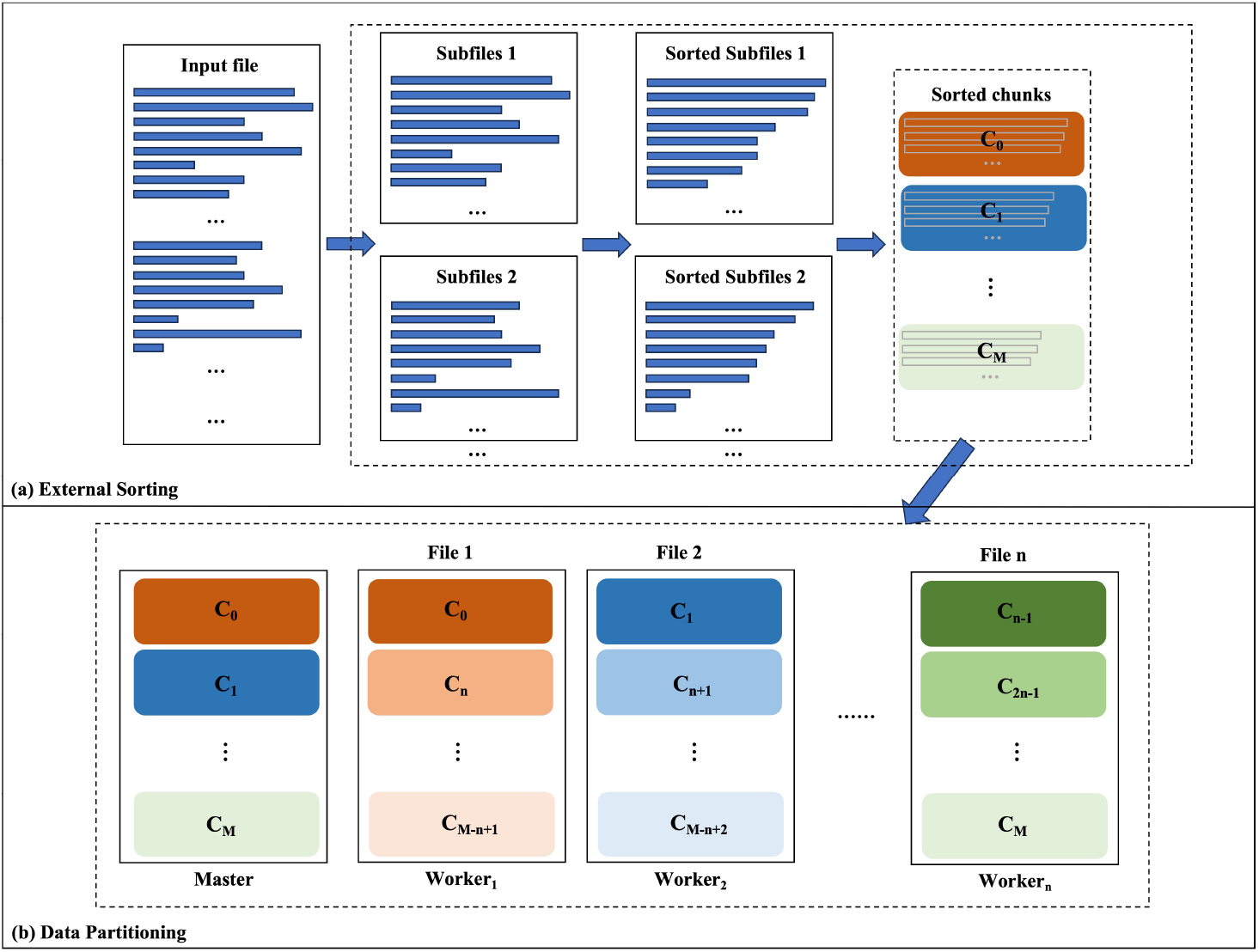
Data partitioning and preprocessing for RabbitCD-HIT: (a) External Sorting, (b) Data Partitioning.

In a greedy clustering algorithm, sequences are processed in sorted order, and each sequence only needs to be compared with those that follow it. As more sequences are absorbed into existing clusters, the remaining comparison workload gradually decreases. If data chunks were distributed sequentially to consumer processes, this asymmetry would lead to significant load imbalance—some processes would quickly finish their tasks while others remain busy.

To mitigate this issue, RabbitCD-HIT adopts a round-robin chunk distribution strategy, as shown in Figure 8(b), cyclically assigning data chunks to consumer processes based on their chunk indices. This strategy provides a balanced static workload distribution across processes during the iterative clustering phase.

### Hierarchical Multi-level Parallelism

RabbitCD-HIT adopts a hierarchical multi-level parallelism strategy across MPI, OpenMP, and vectorization levels. First, MPI-level parallelism distributes sequence chunks and checking workloads across processes through a producer–consumer execution model, together with double-buffered asynchronous pipeline, and streaming processing. Second, OpenMP-level parallelism accelerates different stages on the producer and consumer sides: the producer parallelizes intra-chunk clustering, while the consumers parallelize checking. Third, vectorization-level parallelism accelerates the banded alignment kernel by evaluating multiple dynamic-programming states in parallel using SIMD instructions. These levels work together to improve scalability, memory efficiency, resource utilization, and computational throughput.

#### MPI-Level Parallelism

RabbitCD-HIT reorganizes the chunk-based greedy clustering workflow into an MPI-driven producer–consumer pipeline, as shown in Figure 9.

**Figure 9.**
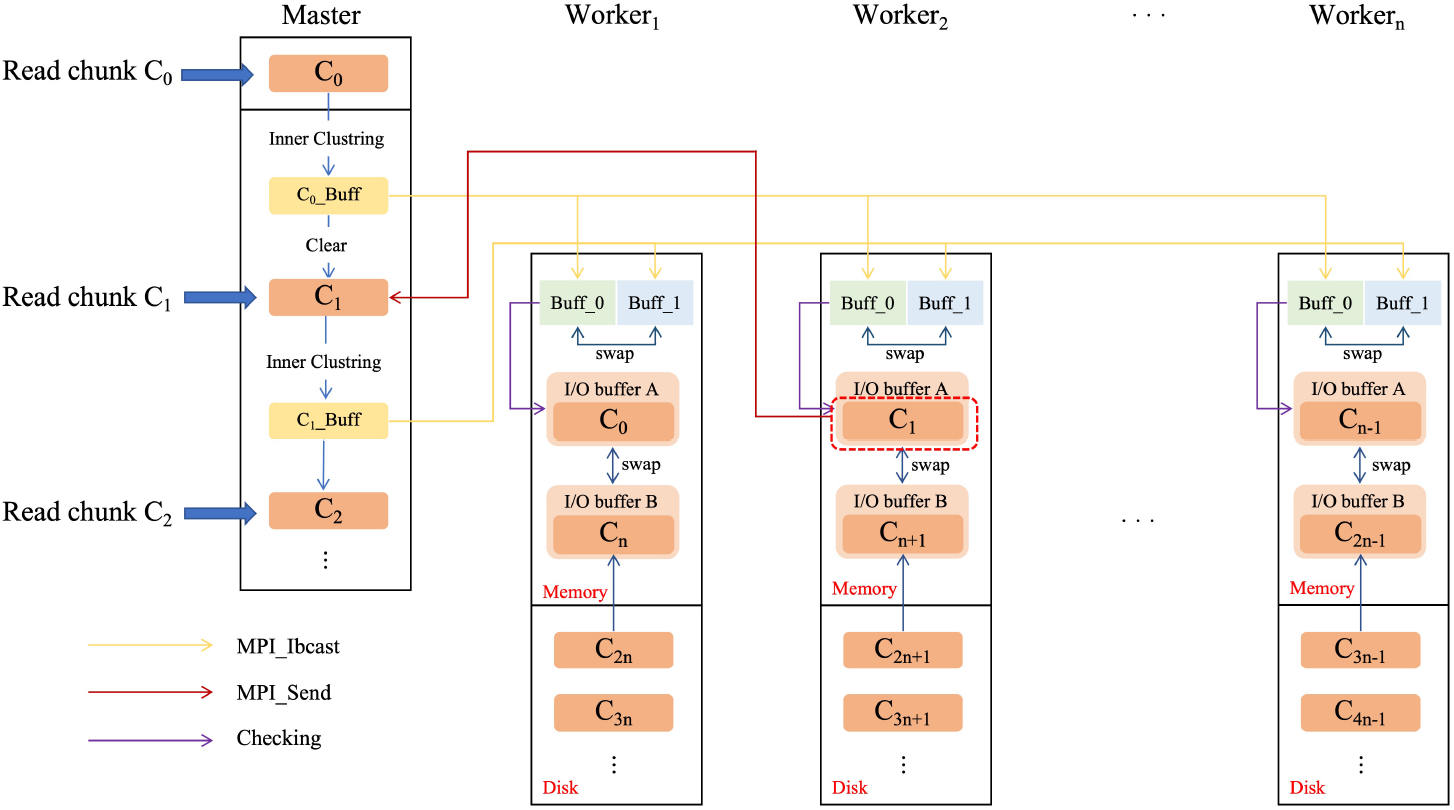
MPI-level producer–consumer pipeline. (i) When the checking of *C*_1_ on *Worker*_2_ is completed, the marking information is immediately sent back to the master process for synchronized state updates, enabling the inner clustering of the next chunk. (ii) Communication buffers are used to store representative information for the current and next chunks; after the current checking round finishes, the communication buffers are swapped and the next round begins. (iii) I/O buffers support on-demand loading during worker-side checking: one buffer holds the current chunk for checking, while the other prefetches the next assigned chunk from disk.

The master process preserves the global greedy order by processing sequence chunks sequentially. For each chunk, the master performs intra-chunk clustering to select representative sequences, encodes their chunk-level indices together with auxiliary metadata, and broadcasts this compact information to all worker processes.

After receiving the broadcast metadata, each worker process decodes the transmitted information, loads the corresponding representative sequences from the externally sorted FASTA files, and constructs a local inverted index from their *k*-mers. It then checks its assigned sequences against the current representatives and generates local clustering results. By broadcasting only representative sequence indices and auxiliary metadata, rather than full sequence data, RabbitCD-HIT substantially reduces MPI communication overhead.

During worker-side checking, the master process loads the next sequence chunk and prepares the necessary data structures for the next round. However, inner clustering of the next chunk cannot be performed immediately, because it depends on the redundancy status updates produced by the checking stage. Specifically, sequences that have already been marked as redundant by previous representatives must be excluded from the selection of new representatives. After receiving and applying these updates, the master performs inner clustering on the next chunk, and broadcasts the representative sequence and metadata information to workers. This iterative workflow continues until all chunks have been processed and the final clustering results have been generated.

After all clustering rounds have been completed, each worker sends its locally stored clustering results to the master process using *MPI Send*. The master aggregates the received results and applies the optimized parallel *IPS*^2^*Ra* sorting method [23] to group sequences belonging to the same cluster. Finally, the clustered results are written to disk in a single batch. Overall, this adaptive streaming workflow enables RabbitCD-HIT to process extremely large protein databases in a memory-efficient manner while maintaining high throughput and avoiding excessive memory consumption.

At the MPI level, RabbitCD-HIT employs a double-buffered asynchronous pipeline to mitigate runtime load imbalance and reduce synchronization overhead. Although the round-robin chunk distribution provides an approximately balanced static workload assignment, it cannot fully eliminate runtime variation among worker processes during the checking stage. Such variation arises because the actual checking cost depends on sequence similarity, filtering effectiveness, and alignment workload. Without such an asynchronous pipeline, faster workers may remain idle after finishing their checking tasks, either waiting for slower workers to complete the current round or waiting for the representative information required for the next round. To address this issue, RabbitCD-HIT adopts a double-buffered asynchronous pipeline, as illustrated in Figure 10. After inner clustering, the master process encodes the representative metadata and initiates a non-blocking broadcast using *MPI Ibcast*. Each worker process maintains two metadata buffers: one is used for the current checking round, while the other receives the metadata for the next round. Thus, while a worker constructs the local inverted index and performs *k*-mer filtering and banded alignment for the current round, the metadata required for the next round can be transferred concurrently. Once the current checking task is completed and the next-round metadata have arrived, the worker swaps the two buffers and proceeds to the next round without waiting for a global synchronization barrier. In addition, clustering results generated during checking are stored locally rather than being collected by the master after every round, further avoiding round-by-round synchronization. By decoupling metadata communication from worker-side checking, the proposed pipeline allows each worker to proceed to the next round as soon as it finishes its current checking task and the next-round metadata are available, without waiting for other workers to reach the same point. Consequently, communication is overlapped with computation, synchronization-induced idle time is reduced, and the impact of uneven checking workloads is mitigated.

**Figure 10.**
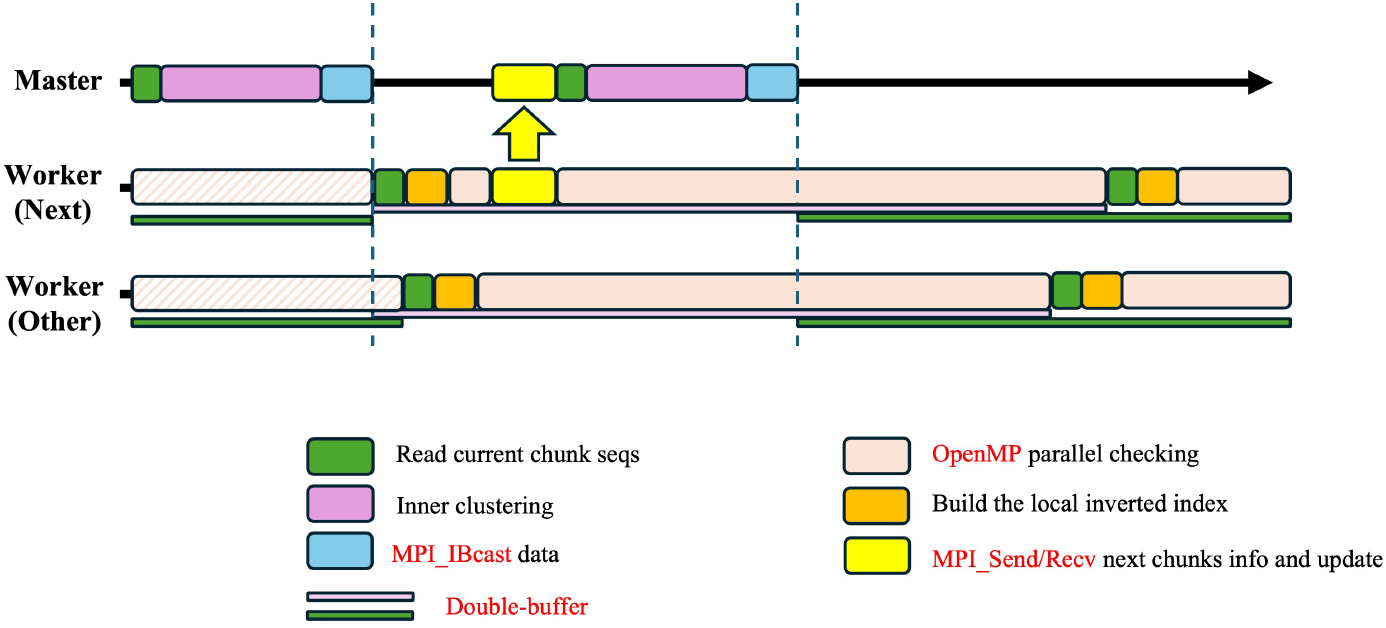
Double-buffered asynchronous pipeline. (i) Data transfer is overlapped with worker-side checking. (ii) Workers use double buffers to receive representative data for the next round while processing the current round, allowing them to proceed asynchronously once the required data are available.

RabbitCD-HIT further introduces a memory-efficient execution scheme to avoid loading the entire sequence database into main memory. Instead, sequence data are loaded on demand from the preprocessed FASTA chunk files. This design bounds memory usage while still allowing disk I/O to be overlapped with communication or computation through prefetching.

On the master side, RabbitCD-HIT processes the database in a chunk-wise manner. In each round, the master loads one sequence chunk, performs inner clustering to identify representative sequences, and encodes the corresponding representative identifiers and auxiliary metadata for broadcasting. After the metadata for the current chunk have been prepared and broadcast to the worker processes, the buffer used for this chunk can be reused to load the next chunk. Therefore, throughout the iterative clustering procedure, the master only needs to maintain a small number of chunk buffers in memory.

On the worker side, RabbitCD-HIT adopts an on-demand loading strategy with double-buffered prefetching during the checking stage. Rather than keeping the entire assigned sub-database in memory, each worker maintains two sequence buffers: one buffer is used for checking the current chunk, while the other prefetches the next required FASTA chunk from disk. Because RabbitCD-HIT processes checking tasks round by round, the next-to-use chunk can be loaded in advance while computation on the current chunk is still ongoing. This strategy reduces memory consumption and hides part of the disk I/O overhead by overlapping data loading with checking computation. It is particularly important for large-scale protein databases, where keeping the full database in memory may exceed the capacity of typical servers.

#### OMP-Level Parallelism

At the OpenMP level, RabbitCD-HIT exploits thread-level parallelism within each MPI process to accelerate both producer-side and consumer-side computation. The hybrid MPI+OpenMP design reduces memory pressure compared with a pure-MPI configuration, because threads within the same process can share sequence buffers, word tables, and other auxiliary data structures instead of maintaining redundant per-process copies. It also reduces the number of MPI communication endpoints and improves intra-node resource utilization, while retaining coarse-grained distributed parallelism across processes. On the master process, OpenMP is used to parallelize inner clustering, thereby preventing the producer from becoming a bottleneck in the distributed pipeline. On the worker processes, OpenMP accelerates the checking stage against the received representative sequences in straightforward manner.

A key objective of RabbitCD-HIT is to maintain a balanced producer-consumer pipeline at the MPI level. Since our distributed framework adopts a single producer process and multiple consumer processes, the throughput of the producer becomes critical at large scale. In particular, if the producer-side inner clustering remains serial, the master process can easily become the bottleneck and prevent the workers from being continuously supplied with new representative information. Therefore, the original single-threaded inner clustering strategy in CD-HIT is not suitable for our producer-consumer design, and an efficient multithreaded inner clustering algorithm is required.

In CD-HIT inner clustering, each query sequence can only be compared against representative sequences that have already been confirmed earlier in the greedy manner, so as to decide whether the query should be retained as a new representative or marked as redundant. This strong sequential dependency makes straightforward parallelization difficult. To overcome this limitation, RabbitCD-HIT redesigns inner clustering as a two-stage lazy-commit multithreaded parallel algorithm. The key idea is to trade a moderate amount of redundant computation for substantially increased parallelism.

In the first stage, RabbitCD-HIT performs parallel candidate generation. All sequences in the current chunk that have not been marked as redundant are temporarily treated as candidate representatives, and their *k*-mers are inserted into a complete inverted index. The entries in each inverted-index bucket are sorted in ascending sequence-ID order. Multiple threads then process different query sequences in parallel. For each query sequence, the thread retrieves the inverted index to collect *k*-mer hits, and performs filtering and alignment to identify sequence pairs that satisfy the similarity threshold. To avoid unnecessary all-to-all comparisons, only smaller-ID sequences are considered as potential candidate representatives for each query sequence, corresponding to the triangular part of the all-to-all comparison space. Since the bucket entries are sorted by sequence ID, the search can stop early once entries with IDs not smaller than the query sequence are reached.

When a sequence pair satisfies the similarity threshold, RabbitCD-HIT records a directed candidate redundancy relation from the query sequence to the smaller-ID sequence. These relations are first stored in thread-local buffers rather than being directly merged into a shared global structure, avoiding contention in the parallel region. After all threads finish candidate generation, the thread-local results are merged for the subsequent commit stage.

In the second stage, RabbitCD-HIT applies a lazy-commit procedure to recover the original greedy incremental clustering behavior. The sequences are scanned sequentially in greedy order. For each sequence, the algorithm checks whether it has a valid candidate redundancy relation to a representative that has already been committed. If such a relation exists, the sequence is marked as redundant; otherwise, it is committed as a new representative. Figure 11 illustrates an example of the commit operation. Initially, sequences 1–5 are all unmarked. Sequence 1 has no valid predecessor and is therefore confirmed as a representative. Sequence 2 points to sequence 1, which has already been committed as a representative, and is thus marked as redundant. Sequence 3 does not satisfy the similarity threshold with any preceding committed representative, so it is retained as a new representative. Sequence 4 points to sequences 2 and 3; however, sequence 2 is itself redundant and therefore cannot serve as a valid representative. Consequently, only sequence 3 remains a valid predecessor for sequence 4, and sequence 4 is marked as redundant with respect to sequence 3. Similarly, sequence 5 points to sequence 4, but this relation becomes invalid once sequence 4 is marked as redundant. As a result, sequence 5 has no valid representative predecessor and is finally committed as a new representative. Although the commit stage is inherently sequential due to the greedy dependency, it only performs lightweight decision updates over the precomputed candidate relations. Therefore, its cost is much smaller than that of the parallel candidate generation stage and has limited impact on the overall performance.

**Figure 11.**
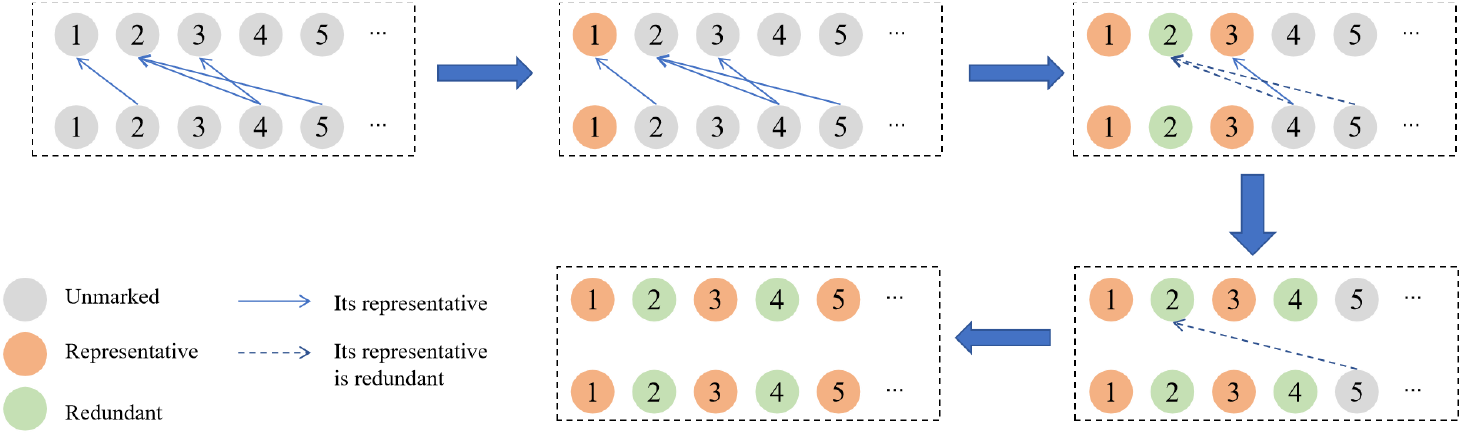
Multithreaded inner-chunk clustering. All unmarked sequences are initially treated as tentative representatives. Threads identify potential redundancy links in parallel by comparing each sequence only with smaller-ID sequences satisfying the similarity threshold. These links indicate possible redundancy rather than final assignments. Final representatives are then committed sequentially in ascending ID order to preserve greedy semantics. A sequence is marked as redundant only if it links to a previously confirmed representative; otherwise, it becomes a new representative.

The proposed two-stage lazy-commit design decouples expensive similarity detection from order-dependent representative confirmation. The computationally intensive *k*-mer filtering and alignment operations are moved to a highly parallel candidate-generation stage, whereas the sequential commit stage only performs lightweight decision updates over the precomputed candidate relations. In this way, RabbitCD-HIT accelerates producer-side inner clustering and helps maintain a balanced producer–consumer pipeline, while preserving the original greedy clustering semantics.

However, when the chunk size is large, the candidate-generation stage may still be expensive. In practice, to achieve high throughput, the chunk size can reach 10^5^ sequences or even larger, making full pairwise candidate generation within an entire chunk time-consuming even with parallel execution and inverted-index filtering. To further reduce this cost, RabbitCD-HIT partitions each chunk into multiple ordered batches and processes these batches sequentially according to the original sequence order. For each batch, candidate redundancy relations are generated in parallel, followed by the lazy-commit procedure to confirm representatives in the original greedy order. Representatives confirmed in earlier batches are immediately incorporated into the representative set and used to filter sequences in later batches. As a result, sequences in later batches that are already covered by earlier representatives can be marked as redundant earlier, reducing unnecessary candidate generation and alignment computations. This ordered-batch strategy provides an additional level of computation reduction on top of the two-stage lazy-commit design, enabling efficient parallel inner clustering while maintaining the correctness of the greedy clustering order.

#### Vectorization-Level Parallelism

At the vectorization level, RabbitCD-HIT optimizes the banded alignment kernel used after the *k*-mer filtering stage. For each query-candidate pair that survives filtering, a diagonal seed is used to determine an alignment band, with the default band width set to 20, and dynamic programming is performed only within this band.

The original CD-HIT alignment kernel uses a banded dynamic-programming procedure with traceback-direction-dependent gap penalties and an additional diagonal-bias score. Given a query sequence *x* and a candidate sequence *y*, only cells (*i, j*) satisfying

,
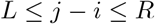

are evaluated, where *L* and *R* denote the left and right boundaries of the selected band. Let *H*_*i,j*_ be the best alignment score at cell (*i, j*) within the band, and let *T*_*i,j*_ ∈ {*D, L, U*} denote its traceback direction, corresponding to diagonal, leftward, and upward transitions, respectively. For each valid cell, the recurrence is computed as

,
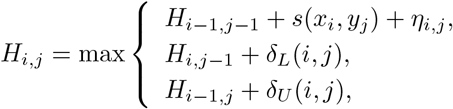

where *s*(*x*_*i*_, *y*_*j*_) is the substitution score between the *i*-th amino acid of *x* and the *j*-th amino acid of *y*. The diagonal-bias term *η*_*i,j*_ is defined as

,
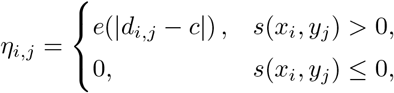

where *d*_*i,j*_ = *j* − *i* is the diagonal index of cell (*i, j*), *c* is the center diagonal of the selected band, and *e*(·) is a small positive bias function. The gap penalties are determined by the traceback direction of the preceding cell:

,
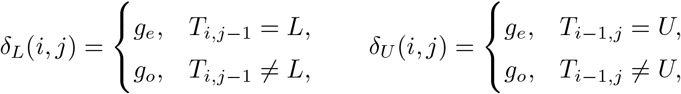

where *g*_*o*_ and *g*_*e*_ denote the gap-opening and gap-extension penalties, respectively. After the maximum score is selected, *T*_*i,j*_ is set to the corresponding transition direction. Boundary cells inside the band are initialized according to the same gappenalty scheme, whereas cells outside the band are treated as invalid and are not evaluated. After all valid cells within the band are evaluated, traceback is performed from the selected terminal cell. The traceback is used to compute the number of identical amino-acid pairs, the alignment length, the alignment distance, and the unmatched terminal regions used in the final clustering decision.

In the original row-major implementation, *H*_*i,j*_ depends on *H*_*i,j−*1_ in the same row, which creates a loop-carried dependency and prevents direct SIMD evaluation of adjacent cells. To remove this dependency, RabbitCD-HIT transforms the banded DP into rotated coordinates:

,
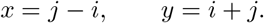

Under this transformation, the three dependency moves become

,
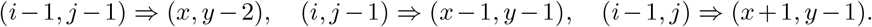

As shown in Figure 12, under this transformation, cells on the same wavefront become independent and the row-wise dependency is removed, enabling multiple cells on a wavefront to be evaluated in parallel using SIMD. However, the parity constraint leaves many inactive positions in the transformed grid, so we compress the rows, making *y*^*′*^ = *y* and 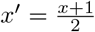. This compression reduces memory overhead, enabling more efficient data access by fitting more active cells into a single vector. It improves cache utilization and optimizes parallel execution, with the wavefront structure intact, allowing independent processing of each wavefront across SIMD lanes.

**Figure 12.**
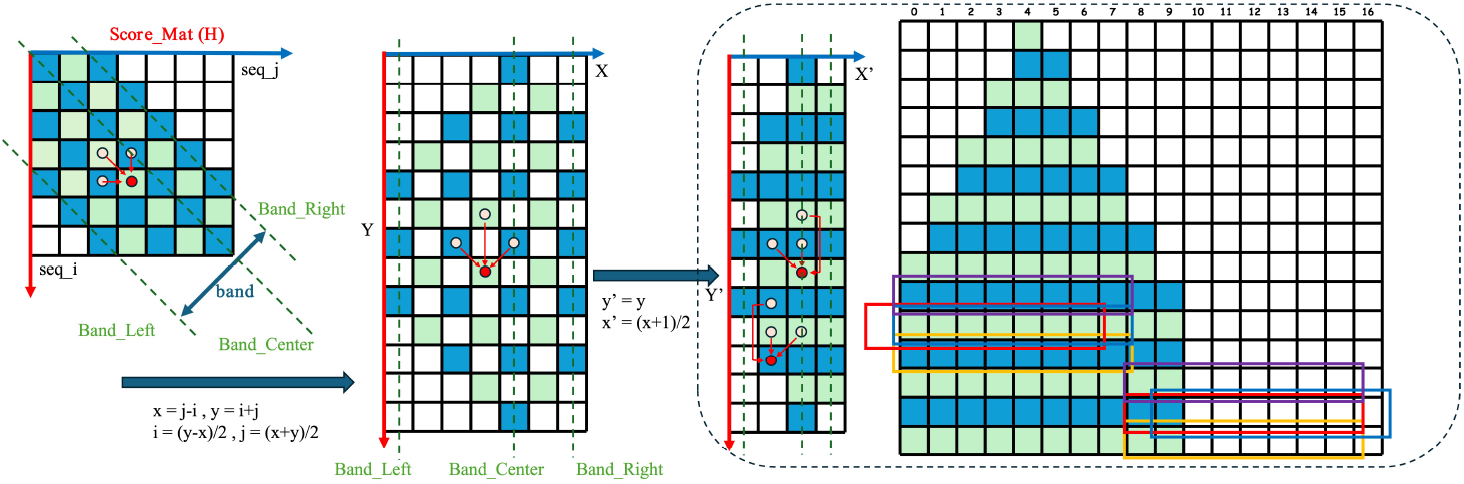
Wavefront banded local alignment in rotated coordinates. By transforming the dynamic programming computation into a wavefront-banded structure, row-wise dependencies are eliminated.

Based on the wavefront formulation described above, we implement a vectorized alignment kernel for AVX-512-based architectures. In this implementation, DP scores are stored as 64-bit integers, allowing eight DP cells on the same wavefront to be evaluated in parallel within a 512-bit vector register. Many highly optimized alignment kernels use shorter integer types to increase SIMD lane utilization. However, RabbitCD-HIT follows the scoring scheme of CD-HIT, where substitution scores and gap penalties are scaled by MAX SEQ (65536). This scaling allows the algorithm to add a small diagonal-bias score when multiple DP cells have the same value, so that the traceback preferentially selects a path closer to the diagonal. As a result, the accumulated DP scores can become too large for shorter integer types, and 64-bit integers are required to preserve numerical correctness and maintain identical behavior with CD-HIT.

### Process-Thread Allocation Strategy

Normally, all MPI processes are assigned the same number of threads, whether they act as the producer or consumers. Since the producer side contains only one process, its performance is directly determined by the number of threads assigned to that process, whereas the consumer side is mainly determined by the total worker-side computing resources.

Improper allocation may cause workload imbalance between the producer and consumers, leading to idle waiting. With too few threads, the master process may become a bottleneck and fail to prepare broadcast data in time; with too many threads, it may finish early and wait for workers to complete checking. Therefore, proper thread allocation is essential for balancing production and consumption and maintaining pipeline efficiency.

To improve the balance between the producer and consumers, RabbitCD-HIT models their execution times under different thread allocations.

Let

- *C*: number of cores (threads) per node;
- *N* : number of nodes;
- *P* : number of MPI processes per node;
- *T* : number of threads per process;
- *chunk seq num*: number of sequences in a chunk;
- *total seq num*: total number of sequences to be clustered;
- *f* (*x*): workload function of producer-side inner clustering for a chunk with *x* sequences;
- *g*(*x*): workload function of consumer-side checking for *x* sequences;
- *α*: single-thread processing rate of the master process for inner clustering;
- *β*: single-thread processing rate of a worker process for representative checking;
- *λ*: empirical correction factor relating the single-thread processing rates of the master and worker processes, where *β* = *λα*. In practice, *λ* is set to a default value based on empirical observations.

Each MPI process is assigned *T* threads, where *T* = *C/P*. The value of *P* is selected by matching the producer-side time *τ*_producer_ and the consumer-side time *τ*_consumer_, with an empirical correction factor *λ* introduced to capture the efficiency difference between inner clustering and checking. The detailed formulation is derived as follows.

To simplify the performance model, we make the following assumptions. First, as clustering proceeds, the effective workloads of both producer-side inner clustering and worker-side checking tend to decrease because more sequences are marked as redundant. We assume that their relative decreasing trends are sufficiently consistent, so a representative pipeline round can be used to estimate the producer-consumer balance. Second, both producer-side and consumer-side computations are assumed to scale linearly with the number of available threads. Third, sequences are assumed to have approximately equal lengths, and all chunks contain the same number of sequences.

Since there is only one producer process, the number of producer threads is

,
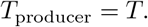

The total number of consumer threads on the worker processes is

,
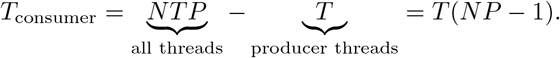

For each round, the producer performs inner clustering on the current chunk (*chunk*_*i*_), while the consumers check the representatives generated from the previous chunk (*chunk*_*i−*1_). The execution times are modeled as

,
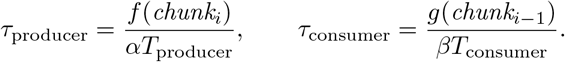

To achieve load balance, we set

,
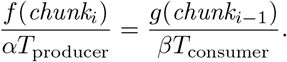

With equal-sized chunks, we use the following coarse-grained workload approximation:

,
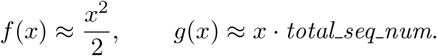

Here, *f* (*x*) approximates the inner chunk candidate space for producer-side inner clustering, while *g*(*x*) approximates checking the representatives from one chunk against the full sequence set. The effects of filtering and alignment are absorbed into the empirical processing rates.

Thus,

,
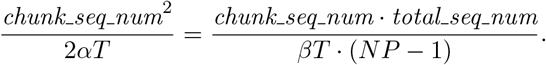

After simplification, we obtain

,
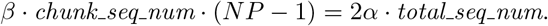

Using *β* = *λα*, the balancing condition becomes

,
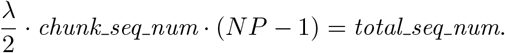

In practice, RabbitCD-HIT iterates over a set of candidate *P* values constrained by the NUMA topology. For each candidate *P*, the corresponding number of threads per process is determined as *T* = *C/P*, and RabbitCD-HIT evaluates how closely the above balance condition is satisfied. When several candidates provide similar balance, the one with a larger *T* is preferred, so that more threads are assigned to the single master process. The final allocation also considers NUMA locality: threads are preferentially placed within the same NUMA domain, and a process may span nearby NUMA domains within the same socket when necessary. This strategy balances producer-side and consumer-side throughput while slightly favoring the master-side computation and preserving good memory locality. The empirical factor *λ* is set according to the observed relative cost between inner clustering and checking.

### Clustering Consistency

To ensure that RabbitCD-HIT preserves the clustering semantics of the original CD-HIT, we compare the outputs of the two programs under both the default mode and a strictly aligned mode.

In the default mode, both RabbitCD-HIT and CD-HIT use the standard fast greedy assignment strategy. Under this strategy, sequences are processed in the greedy order, and each query sequence is assigned to the first representative sequence that satisfies the similarity threshold. Under the default mode, RabbitCD-HIT produces exactly the same representative sequence set as the original CD-HIT. However, complete cluster assignment may still be affected by the traversal order of candidate representatives and the handling of equal-length sequences, leading to minor inconsistencies in the output .clstr files. Since the representative sequence set is identical, these inconsistencies have limited impact on clustering quality.

Furthermore, we use a strictly assign mode (choosing the best hit instead of the first hit) to compare the full clustering assignments. In this mode, both RabbitCD-HIT and CD-HIT are executed with -g 1, so that each sequence is assigned to the most similar eligible representative rather than the first qualified one. RabbitCD-HIT is additionally executed with -fo 1, which enforces the same processing order for equal-length sequences as CD-HIT. When multiple representatives have the same highest similarity to a query sequence, both implementations assign the query to the representative with the smaller sequence ID. Under the strictly aligned mode, all sequence-to-representative pairs produced by RabbitCD-HIT and CD-HIT are exactly identical, confirming that RabbitCD-HIT preserves CD-HIT-compatible clustering semantics.

## Acknowledgements

We sincerely thank Weizhong Li, the original developer of CD-HIT, for his valuable support and helpful suggestions during this study.

## Availability of data and materials

The 90% non-redundant protein sequence database generated by RabbitCD-HIT, including the representative sequence set and the complete clustering results, is publicly available at: https://1drv.ms/f/c/d96807d248ed9ca3/IgB-EvcyFSedTJ3rrKaRTpqBAZbb_SKnaIXXBnN-eP_lI8U?e=z6H6beRabbitCD-HIT is freely available from https://github.com/RabbitBio/RabbitCDHIT.

RabbitCD-HIT is written in C++ and has been tested on 64-bit Linux Systems.

The experimental datasets used in this study, including subsets containing 2, 10, 50, and 100 million sequences, are available from Zenodo under DOI: https://doi.org/10.5281/zenodo.19812052.

## Authors’ contributions

Z.Y. and W.L. designed and supervised this study. Y.G. and G.M. codesigned this study, developed the software, and performed the analyses. L.Y., Y.Y., and Tong Z. helped to perform the experiment analyses. Tiancheng Z., Z.L., X.D., B.N., W.L., and B.S. performed the analyses. Y.G., G.M., Z.Y., W.L., and B.S. wrote the manuscript. All authors read and approved the final manuscript.

## Declarations

### Ethics approval and consent to participate

Not applicable.

### Consent for publication

Not applicable.

